# Magnesium-28: A Novel Self-Theranostic Approach through Targeting Metabolic Enzyme Disruption and Intracellular Radiation

**DOI:** 10.1101/2025.05.12.653584

**Authors:** Tran Van Luyen

## Abstract

The limitations of conventional cancer therapies, such as low selectivity and significant side effects, necessitate innovative approaches. This study proposes a pioneering self-theranostic strategy using Magnesium-28 (Mg-28) alone, enabling simultaneous diagnosis, therapy, and therapy control. Exploiting the elevated Mg2+ demand in cancer cells, Mg-28 self-targets Mg-dependent enzymes (e.g., DNA/RNA polymerases, hexokinase, telomerase) within intracellular organelles like the nucleus and mitochondria, without bio-chemical carriers or nanoparticles as recently methods. A theoretical model, based on the Mg-uptake coefficient, predicts selective Mg-28 accumulation in tumors following intravenous administration. The Mg-28 chain decays into Aluminum-28 then Silicon-28, delivering highly localized irradiation via beta particles, Auger electrons, and recoil ions to critical intracellular structures, while disrupting essential Mg-dependent enzymes for a dual mechanism of radiotherapy and multi-enzyme inactivation. Simulations of Linear Energy Transfer (LET), radiation range, and absorbed dose show that nanogram-scale amounts of Mg-28 can deliver 60-400 Gy to tumors ranging from 0.03 mg to 500 g in size, suggesting potential cytotoxicity that could be effective across not only a broad range of stages but also types of cancer due to the fundamental role of magnesium in cancer cell metabolism and proliferation. Mg-28 and its daughter’s gamma emissions support early tumor detection and real-time treatment monitoring, enhancing precision. As the first proposed single-isotope theranostic approach leveraging magnesium dependency, this innovative strategy provides a robust foundation for future preclinical and clinical investigations aimed at validating its therapeutic efficacy, pharmacokinetics, and biosafety inaugurating a novel hypothesis for cancer therapy.

## I. Introduction

Cancer remains a paramount global health challenge, responsible for approximately 10 million deaths worldwide in 2020 [1]. Despite significant progress in medical oncology, conventional treatment modalities such as surgery, chemotherapy, radiotherapy, and immunotherapy are often limited by reduced efficacy in advanced stages, debilitating systemic side effects, and the frustrating issue of tumor relapse due to the incomplete eradication of cancer cells [2].

The advent of targeted therapies, including nanoparticle delivery systems [3], monoclonal antibodies [4], peptides [5], CRISPR-Cas9 [6], and CAR-T cells [7], has offered improved precision in targeting cancer cells. However, these approaches predominantly focus on extracellular targets and often require complex delivery mechanisms. A critical gap persists in our ability to effectively target key intracellular processes, particularly the enzymes essential for cancer cell metabolism and replication.

While radioisotopes like Iodine-131, Phosphorus-32, Lutetium-177, Holmium-166, and Yttrium-90 have advanced the field of extracorporeal radiotherapy, including techniques like brachytherapy [8, 9], these methods typically rely on carriers to deliver isotopes to the vicinity of tumors and primarily exert their effects extracellularly, thus limiting their impact on intracellular cancer cell machinery [10]. For instance, Iodine-131, while effective for thyroid cancer, carries the risk of secondary malignancies [11].

A fundamental challenge in current cancer treatment is the failure to comprehensively disrupt the core processes that drive cancer progression: uncontrolled proliferation, limitless replicative potential (immortality), invasion, metastasis, and the sustained energy supply required for these processes. Intriguingly, all these critical functions are heavily reliant on magnesium (Mg²⁺) as an essential cofactor for numerous Mg-dependent enzymes [12]. Tumor cells, characterized by their rapid proliferation and heightened metabolic demands, exhibit a significantly elevated requirement for Mg²⁺ compared to normal cells. This metabolic vulnerability presents a unique and underexplored therapeutic opportunity.

Historically, Magnesium-28 (Mg-28) has been utilized as a valuable tracer in metabolic studies, particularly in plant biology and in investigating the pathophysiology of diabetes [13–15]. However, its potential as a therapeutic agent in the direct treatment of cancer remains largely unexplored. This study introduces Mg-28 as a dual-action agent that combines targeted intracellular irradiation with the direct inactivation of Mg-dependent enzymes. A potential concern regarding the use of Mg-28 stems from the fact that Mg²⁺ is a cofactor for over 300 enzymes, raising questions about potential off-target effects in vital organs such as the lungs, liver, and brain. However, this research demonstrates the contrary: Mg-28 exhibits a strong propensity to selectively concentrate on malignant tumors, the very sites where cell survival, proliferation, invasion, and metastasis necessitate a magnesium demand far exceeding that of healthy tissues. Following intravenous administration, Mg-28 is hypothesized to selectively accumulate within these tumor cells, competitively replacing stable Mg²⁺ in crucial enzymes. The subsequent decay of Mg-28 emits beta particles, Auger electrons, and recoil ions, directly targeting intracellular structures such as the nucleus and mitochondria. This dual mechanism aims to overcome resistance, minimize off-target side effects, and enable self-theranostic applications, integrating both therapy and diagnostics within a single agent.

The overarching objective of this work is to comprehensively analyze the Mg-28 approach, demonstrating its feasibility as a groundbreaking strategy that combines the precision of targeted intracellular chemotherapy with radiotherapy, while minimizing damage to healthy tissues and enabling early diagnosis and dynamic monitoring of therapeutic progress. This research seeks to unlock the transformative potential of Mg-28 as a next-generation theranostic platform in the ongoing fight against cancer. In fact, this approach could be a fundametal for preclinical and clinical evaluation in the future.

## II. Methodology

This theoretical study evaluates the feasibility of Mg-28 as a precision cancer therapy through a comprehensive analysis encompassing five key aspects: (1) the fundamental principles of metalloenzyme inactivation, (2) the mechanism of Mg-dependent multienzyme disruption due to changes in cofactor valence and ionic radius, (3) the calculation of linear energy transfer (LET) and the range of emitted radiation particles, including recoil effects, (4) the quantification of absorbed dose in tumors of varying volumes and the assessment of systemic dose, and (5) the tumor-specific uptake of Mg ions driven by the metabolic demands of cancer cells. This approach builds upon the basic principles of metalloenzyme inactivation detailed in our previous publications [16, 17].

### II.1. Principles of Metalloenzyme Inactivation

The concept of substituting stable metal ion cofactors in metalloenzymes with suitable radioisotopes to induce enzyme inactivation forms the basis of this approach. Ideal radioisotopes for this purpose emit beta particles or Auger electrons and possess a half-life that is neither excessively long (to minimize prolonged patient radiation exposure) nor too short (to ensure practical clinical application). Furthermore, their decay products should be isotopes of elements that do not function as cofactors at the enzyme’s active site. Magnesium-28 (T₁/₂ ≈ 21 h) meets these criteria for targeting Mg-dependent enzymes, decaying into Al-28 and subsequently Si-28, with emissions capable of disrupting the function of these critical enzymes in cancer cells [18].

### II.2. Mechanism of Mg-Dependent Enzyme Inactivation

Magnesium ions (Mg²⁺) typically stabilize the active sites of Mg-dependent enzymes by forming six-coordinate bonds with oxygen atoms from carboxylate and phosphate groups [19–22]. This coordination is essential for substrate binding and catalytic activity. The radioactive decay of Mg-28 into Aluminum (Al³⁺) and then Silicon (Si⁴⁺) results in ions with higher charges and smaller ionic radii (Al³⁺: 0.50 Å; Si⁴⁺: 0.40 Å) compared to Mg²⁺ (0.72 Å) [23, 24]. This substitution disrupts the delicate electrostatic interactions within the active site, leading to structural stress, distortion, weakened substrate binding, and ultimately, impaired or abolished catalytic efficiency. Additionally, the recoil of Al-28 and Si-28 ions (with a displacement of 0.022–1.5 Å) can cause local distortions at the enzyme active site, further contributing to the loss of function given the precise spatial requirements for enzymatic catalysis. The high LET particles (beta particles and Auger electrons) emitted during decay also contribute to enzyme inactivation by breaking covalent and non-covalent bonds within the apoenzyme and generating free radicals that can denature surrounding proteins.

### II.3. LET and Radiation Range

The linear energy transfer (LET) and the range of beta particles, Auger electrons, and recoil ions emitted during Mg-28 decay were calculated using the NIST E-STAR program [25] and MIRD (Medical Internal Radiation Dose) data [26, 27]. For recoil ions (Al-28 and Si-28), recoil energies were derived based on the principle of conservation of momentum following beta particle emission from the parent Mg-28 nucleus:

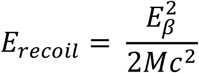

Where: E*_recoil_*is the recoil energy of the daughter nucleus (Al-28 or Si-28),

E_β_ is the energy of the emitted beta particle, and M is the mass of the daughter nucleus.

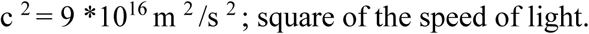

These calculated energy values were then used to model the LET and range of these recoil ions in different tissue types.

### II.4. Absorbed Dose Calculations

Absorbed dose calculations were performed for tumors of different volumes (T_0_–T_5_) and for the whole body of a 60-kg individual using the MIRD program [28] and open-access nuclear data [26, 27]. Calculations considered the administration of 0.1 to 6.2 ng of Mg-28 and modeled three delivery scenarios: intravenous injection with no tumor-specific uptake, intravenous injection with high tumor-specific uptake (based on the Mg uptake coefficient), and direct injection into the tumor. Additionally, absorbed doses were evaluated for different treatment regimens involving 62, 300, and 400 Mg-28 ions per cell, corresponding to varying degrees of inactivation of approximately 300 Mg-dependent enzymes.

### II.5. Tumor-Specific Mg Uptake

The selective accumulation of magnesium ions (Mg²⁺) by cancer cells, compared to their normal counterparts, forms the fundamental basis for employing the radioisotope Mg-28 in targeted cancer therapy. This differential uptake arises primarily from the significantly higher replication rates of cancerous tissues, leading to an increased demand for Mg²⁺, a crucial cofactor for numerous enzymes involved in DNA replication, protein synthesis, and energy metabolism [12, 21]. While intracellular concentrations of stable Mg ions may be comparable between normal and cancerous cells, the dynamic process of cell division creates a substantial disparity in the overall Mg uptake at the tissue level.

To quantify this difference, we model the reproductive capacity of healthy and cancerous tissues over time. Let A and B represent the number of healthy and cancer cells, respectively, after time t, with na and nb being the number of doubling periods, and Ta and Tb representing the replication cycle times for healthy and cancer cells. The proliferation can be expressed as:

*Healthy tissue:*

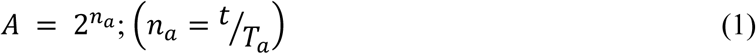

*Cancer tissue:*

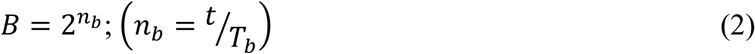

Where:

- A and B: Number of healthy cells and cancer cells respectively.
- n_a_ and n_b_: Number of doubling periods of healthy tissue and cancerous tissue.
- t: Actual copy time.
- T_a_ and T_b_: The replication cycle of healthy cells and cancer cells.

The growth ratio between cancerous tissue and healthy tissue is expressed as follows:

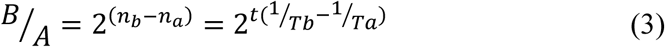

Introducing the doubling time ratio k = Ta/Tb. This factor highlights differences in cell division dynamics. Unlike normal cells, cancer cells are not regulated by Cyclin-Dependent Kinases (CDKs) [29] which ensure genomic integrity, accurate protein synthesis, and complete DNA repair in healthy cells, thus prolonging their replication time. For this reason, k factor is always bigger than one (k>1).

Equation (3) becomes:

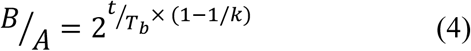

Rapid replication in cancer cells creates a disproportionately high demand for resources essential for survival and division, including Mg ions. This demand doubles during the M phase of the cell cycle. Therefore, this ratio, when normalized to the initial number of cells known as the Mg-28 uptake coefficient. This preferential Mg uptake by cancer cells is the cornerstone of the Mg-28 therapy. The elevated demand for Mg²⁺ in rapidly dividing cancer cells acts as a natural driving force for the selective accumulation of the Mg-28 radioisotope within the tumor microenvironment. This intrinsic targeting mechanism eliminates the need for complex biochemical carriers or nanoparticles, simplifying the treatment process and reducing potential off-target toxicities. The high Mg-uptake coefficient not only enhances the intracellular delivery of Mg-28 for enzyme inactivation and irradiation but also underpins its potential for early diagnosis and real-time monitoring, as even small tumors exhibit a measurable increase in Mg accumulation. This coefficient is also the basis for calculating absorbed doses and enzyme inactivation in intravenous treatment regimens due to involves a mechanism of energy transfer from Mg-28 decay to microenvironment inside cancer cells cause the moleccular bonds break.

The rapid proliferation of cancer cells generates an exceptionally high demand for vital resources, including magnesium ions (Mg²⁺), which doubles during the M phase of the cell cycle. This increased requirement forms the basis of the Mg-28 uptake coefficient, a ratio normalized to the initial number of cells. This preferential uptake of Mg²⁺ by cancerous cells is the fundamental principle behind Mg-28 therapy. The heightened need for Mg²⁺ in rapidly dividing cancer cells acts as an inherent driving force, enabling the selective accumulation of the Mg-28 radioisotope within the tumor microenvironment. This intrinsic targeting mechanism obviates the need for complex biochemical carriers or nanoparticles, thereby simplifying the treatment process and potentially reducing off-target toxicities. The high Mg-uptake coefficient not only enhances the intracellular delivery of Mg-28 for enzyme inactivation and irradiation but also supports its potential for early diagnosis and real-time monitoring, as even small tumors exhibit a detectable increase in Mg accumulation. Furthermore, this coefficient serves as the foundation for calculating absorbed doses and enzyme inactivation in intravenous treatment regimens, where the decay of Mg-28 and subsequent energy transfer within the cancer cell microenvironment led to the disruption of molecular bonds.

## III. Results

### III.1. Mg-Uptake Coefficient

The Mg-uptake coefficient (B/A), a key determinant of Mg-28 distribution, caculated by the equation (4) with an absumption k = 2, demonstrates a significant increase with tumor size and the number of replication cycles.

As demonstrated in Table 1, this coefficient increases dramatically with the number of replication cycles (n_b_) and the value of k. For instance, even with a modest k value of 2, the Mg-uptake coefficient escalates from 1.8×10^2^ at 15 cycles (T_0_ tumor) to 7.1×10^5^ at 39 cycles (T_5_ tumor). This highlights the profound ability of growing tumors to selectively accumulate Mg ions. This trend reflects the elevated magnesium demand of rapidly proliferating cancer cells, which enhances the selective targeting of Mg-28 to larger and more metabolically active tumors compared to smaller or less active ones, and significantly more than normal cells.

**Table 1:**
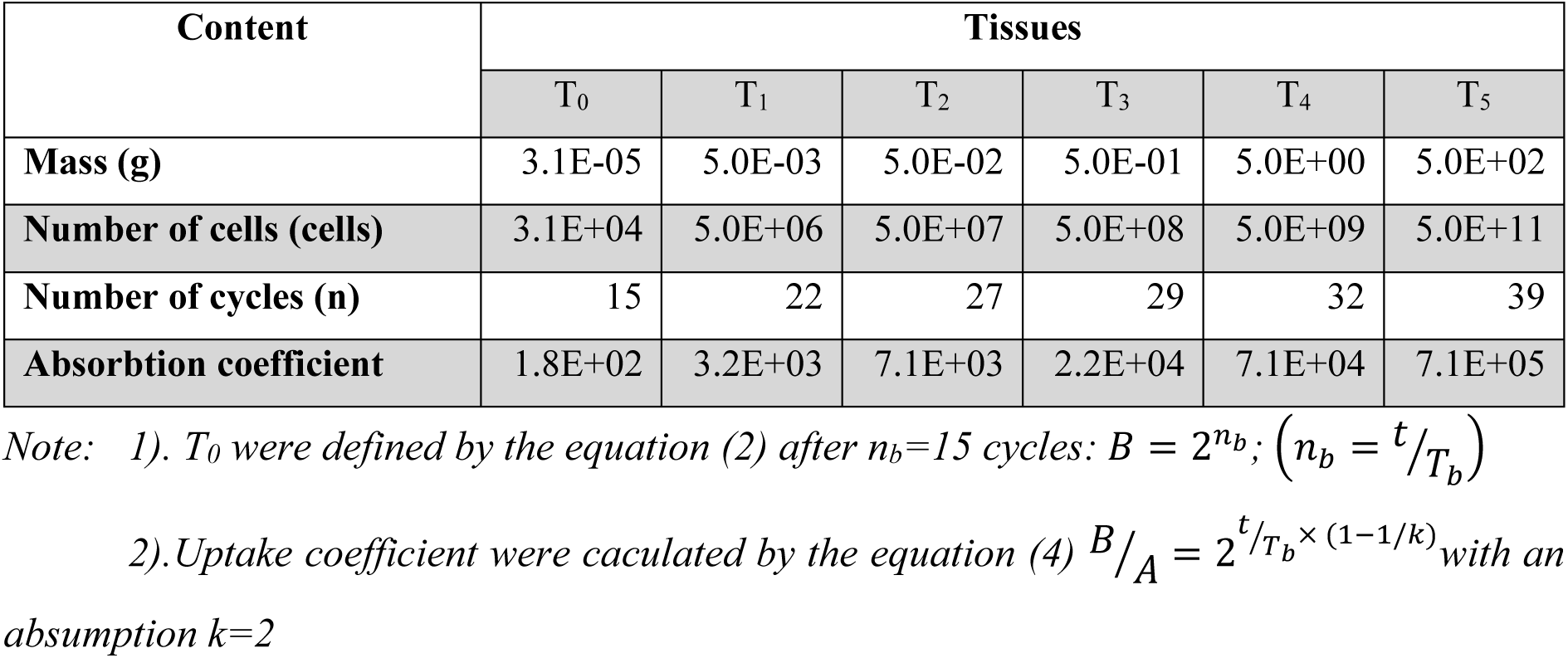
Tissue data, Mg-28 absorbtion coefficient (k = 2)

### III.2. LET and Particle Range

The Linear Energy Transfer (LET) values and corresponding ranges for the particles emitted during Mg-28 decay are presented in Table 2.

**Table 2.**
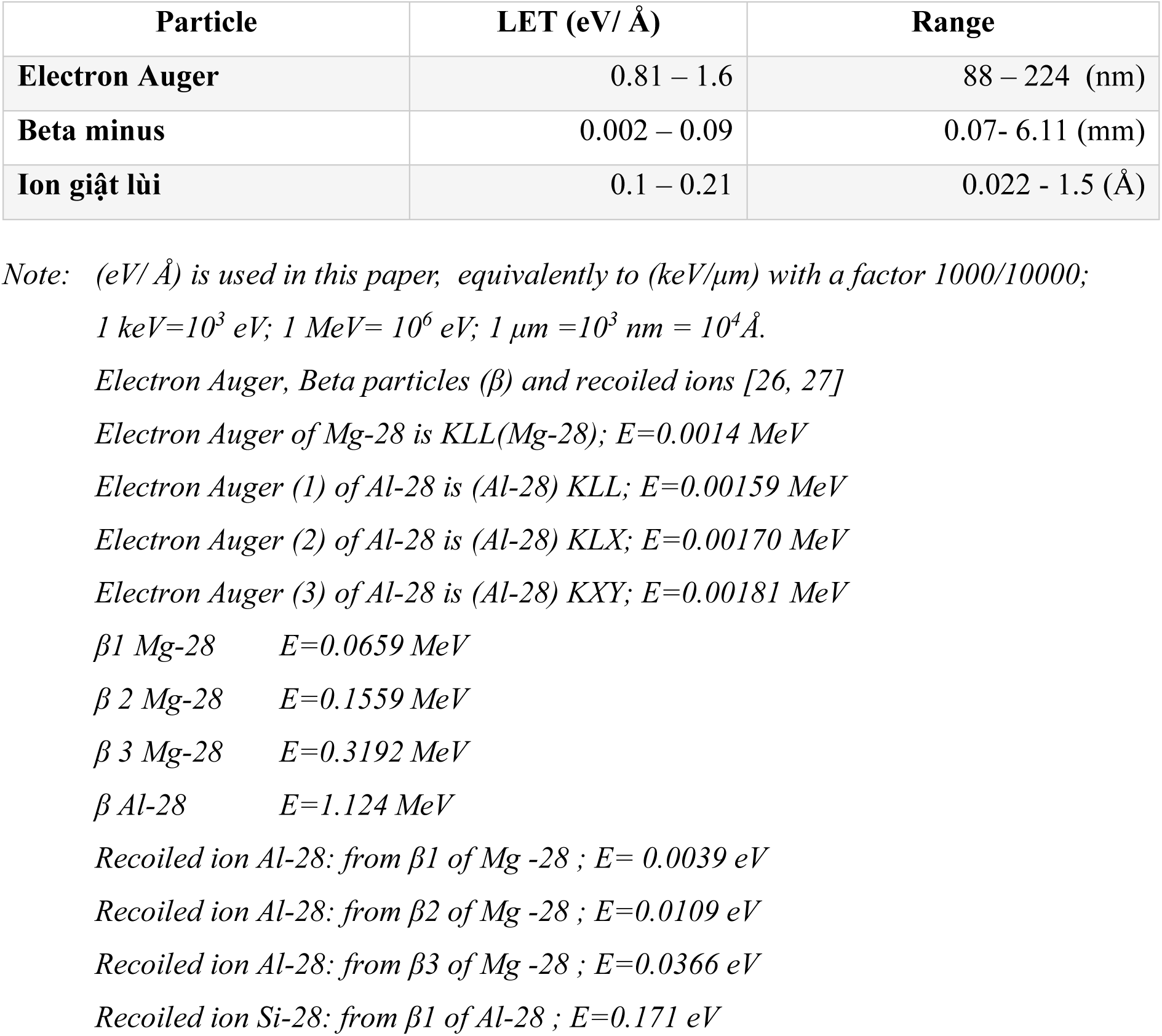
LET and range of electron Auger, beta minus and recoil ion.

Beta-minus particles exhibit LET values ranging from 0.002 to 0.09 eV/Å with a range of 0.07 to 6.11 mm. Auger electrons demonstrate higher LET values, ranging from 0.81 to 1.6 eV/Å, but with a shorter range of 88 to 224 nm. Recoil ions (Al-28 and Si-28) have LET values between 0.1 and 0.21 eV/Å and a very short range of 0.022 to 1.5 Å. These values (Table 3) indicate that while Auger electrons possess the highest LET, recoil ions, despite their limited range, can effectively disrupt enzyme bonds due to their focused action at the molecular level.

**Table 3:** Impact of LET on the biological molercular bonds.

| Bonds [30] | Length (Å) | Energy bond (kJ/mol) | Beta minus (kJ/mol) | Auger electron (kJ/mol) | Recoiled ions (kJ/mol) |
| --- | --- | --- | --- | --- | --- |
| Van der Waals | 3.0-4.0 | ~ 2.4 - 4 | **7.08 - 34.84** | **311.81 - 612.41** | **38.12 - 81.08** |
| Hydro (H-bond) | 1.5-2.0 | ~4 - 40 | 3.54 - **17.42** | **155.91 - 306.21** | 19.06 - **40.54** |
| P–O (in ATP/ADP) * | ~1.60 | ~30.5 | 2.83 - 13.94 | **124.72 - 244.96** | 15.25 - **32.43** |
| Ion-ion (Coulombic) | 2.0-4.0 | ~20 - 200 | 7.08 - 34.84 | **311.81 - 612.41** | 38.12 - 81.08 |
| N–N (Nitrogen-Nitrogen) | ~1.45 | ~163 | 2.57 - 12.63 | 112.88 - **222.00** | 13.82- 29.39 |
| S–S (Disulfide) | 2.05 | ~250 | 3.64 - 17.82 | 156.31 - **381.20** | 19.46 - 41.54 |

Note: \* Energy released when ATP hydrolyzes to ADP; LET: Linear Energy Transfer.

### III.3. Absorbed Dose

Absorbed dose calculations by MIRD code [28], using 0.1 ng of Mg-28 administered intravenously over 21 hours (Table 4), showed the potential for highly effective tumor killing with minimal impact on healthy tissues. Three regimens: (a) whole-body cell dose, (b) Mg-uptake-coefficient cell dose, and (c) direct-tumor cell dose showed significantly different results. In regimen (a), the doses ranged from E-11 to E-04 Gy; in regimen (b), the doses increased from E-09 to E+02 Gy; and in regimen (c), the doses decreased from E+04 to E-02 Gy in the corresponding T_0_-T_5_ tumors (0.03 mg – 500 g). This indicates that the Mg-uptake coefficient significantly increased the cell dose compared to the whole-body distribution.

**Table 4.** Absorbed dose (Gy) impact to tumor of different regimens and amount of Mg-28 required.

| Content | Cancer type |  |  |  |  |  |
| --- | --- | --- | --- | --- | --- | --- |
|  | T<sub>0</sub> | T<sub>1</sub> | T<sub>2</sub> | T<sub>3</sub> | T<sub>4</sub> | T<sub>5</sub> |
| Number of cells | 3,28E+04 | 5,00E+06 | 5,00E+07 | 5,00E+08 | 5,00E+09 | 5,00E+11 |
| Intravenous cell dose distributed uniformly whole body (Gy)-a | 1,01E-11 | 1,55E-09 | 1,55E-08 | 1,55E-07 | 1,55E-06 | 1,55E-04 |
| Intravenous cell dose with Mg absorption coefficient (Gy)-b | 1,83E-09 | 3,46E-06 | 1,09E-04 | 3,46E-03 | 1,09E-01 | 1,09E+02 |
| Cell dose injected directly to tumor (Gy)-c | 3,04E+04 | 2,23E+02 | 2,23E+01 | 2,23E+00 | 2,23E-01 | 2,23E-02 |
| Tumor dose -a (Gy) | 3,31E-07 | 7,75E-03 | 7,75E-01 | 7,75E+01 | 7,75E+03 | 7,75E+07 |
| Tumor dose -b (Gy) | 6,00E-05 | 11,15E+00 | 11,15E+03 | 11,15E+05 | 11,15E+08 | 11,15E+13 |
| Tumor dose -c (Gy) | 9,97E+08 | 38,75E+08 | 38,75E+08 | 38,75E+08 | 38,75E+08 | 38,75E+09 |
| Tumor dose - d (Gy) | 6,01E+01 | 6,01E+01 | 6,01E+01 | 6,01E+01 | 6,01E+01 | 6,01E+01 |
| Tumor dose - e (Gy) | 3,12E+02 | 3,12E+02 | 3,12E+02 | 3,12E+02 | 3,12E+02 | 3,12E+02 |
| Tumor dose - f (Gy) | 4,15E+02 | 4,15E+02 | 4,15E+02 | 4,15E+02 | 4,15E+02 | 4,15E+02 |
| Whole body absorbed dose case d (Gy) | 3,52E-01 | 2,03E-02 | 8,91E-03 | 2,82E-03 | 8,91E-04 | 8,91E-05 |
| Whole body absorbed dose case e (Gy) | 1,72E+00 | 9,81E-02 | 4,42E-02 | 1,44E-02 | 4,42E-03 | 4,42E-04 |
| Whole body absorbed dose case f (Gy) | 2,33E+00 | 1,32E-01 | 5,92E-02 | 1,92E-02 | 5,92E-03 | 5,92E-04 |
| Amount of Mg-28 required in case d (ng) | 9,31E-01 | 5,33E-02 | 2,42E-02 | 7,51E-03 | 2,44E-03 | 2,44E-04 |
| Amount of Mg-28 required in case e (ng) | 4,62E+00 | 2,63E-01 | 1,22E-01 | 3,71E-02 | 1,22E-02 | 1,22E-03 |
| Amount of Mg-28 required in case f (ng) | 6,21E+00 | 3,51E-01 | 1,63E-01 | 5,04E-02 | 1,63E-02 | 1,63E-03 |

*Note: The absorbed dose of Mg-28 were calculated during 21 hours.*

Based on these results, we proposed intravenous regimens (d), (e), (f) with 62, 300, and 400 Mg-28 ions per cell, respectively, capable of competing for 20%, 100%, and 133% of the total 300 Mg-dependent enzymes, and obtained the following results: The doses ranging from 60 to 415 Gy were achieved on T_0_ to T_5_ tumors with a relatively low maximum amount of only 6.2 ng of Mg-28. Notably, the highest effective systemic absorbed dose among the model regimens remained below 2.4 Gy, indicating a favorable safety profile. These results demonstrate that the absorbed dose delivered by Mg-28 is sufficient to induce significant cytotoxicity in cancer cells across a wide range of tumor sizes without causing significant damage to surrounding healthy tissues and with minimal radioisotope exposure.

## IV. Discussion

### IV.1. Tumor Growth, Replication Cycle, and Magnesium Uptake

Cancer cell replication cycles are known to vary depending on tumor type and microenvironmental conditions. Our data, presented in Table 1, indicate a clear relationship between tumor size, the number of replication cycles, and the Mg-uptake coefficient. For instance, a microscopic tumor at stage T_0_ (mass ∼3.1×10^−5^ g, approximately 3.1×10^4^ cells) after 15 replication cycles exhibits an Mg-uptake coefficient of 1.8×10^2^. As the tumor progresses through stages T_1_ to T_5_, with increasing mass and number of replication cycles, the Mg-uptake coefficient demonstrates a significant upward trend, reaching 7.1×10^5^ for a large T_5_ tumor (mass ∼500 g, ∼5.0×10^11^ cells) after 39 cycles, assuming a replication cycle ratio (k) of 2 between cancerous and healthy tissue.

This substantial increase in the Mg-uptake coefficient with tumor growth underscores the elevated magnesium demand of rapidly proliferating cancer cells. As tumors expand, their metabolic and replicative requirements for magnesium, a crucial cofactor for numerous enzymes involved in these processes, escalate. Consequently, Mg-28 is preferentially accumulated in larger and more aggressive tumors, making it an increasingly effective agent for targeted radiotherapy in advanced stages.

Furthermore, the relatively high Mg-uptake coefficient observed even in early-stage T_0_ tumors suggests a promising avenue for early cancer detection, which will be discussed further in the subsequent section. The dependence of the Mg-uptake coefficient on the number of replication cycles and the k-factor highlights the importance of considering tumor growth dynamics in optimizing Mg-28-based treatment strategies. Tumors with faster replication rates (higher k values) are expected to exhibit even greater Mg-28 accumulation, potentially enhancing therapeutic efficacy and reducing the required dosage.

### IV.2. Early Diagnosis and Real-Time Monitoring

The exceptionally high Mg-uptake coefficient observed even in early-stage, microscopic tumors (T_0_: 1.8×10^2^, as shown in Table 1) provides a strong foundation for early cancer diagnosis. This preferential accumulation of Mg-28 in nascent tumors allows for robust imaging signals via gamma rays, Bremsstrahlung, and X-rays, enabling detection through PET or SPECT imaging. The ability to visualize tumors as small as ∼3.1×10^−5^ g (approximately 31,000 cells) represents a significant advancement over many current diagnostic modalities that may only detect larger tumor masses.

Moreover, the continuous uptake of Mg-28 by tumor cells, sustained throughout the treatment period, facilitates real-time monitoring of therapeutic response. Changes in Mg-28 concentration within the tumor, as visualized through imaging, can provide immediate feedback on treatment efficacy, allowing for timely adjustments to the therapeutic regimen. This dual diagnostic and therapeutic (theranostic) functionality, achieved with a single isotope, simplifies the clinical workflow and eliminates the need for paired diagnostic and therapeutic agents, representing a key advantage of the Mg-28 method.

### IV.3. Enzyme Inactivation and Intracellular Irradiation

Data from Table – 2 reveals the Linear Energy Transfer (LET) and range of the particles emitted during Mg-28 decay. Auger electrons exhibit the highest LET (0.81 – 1.6 eV/Å), followed by recoil ions (0.1 – 0.21 eV/Å), while beta-minus particles have a lower LET (0.002 – 0.09 eV/Å) but a much longer range (0.07 – 6.11 mm).

The high LET of Auger electrons and recoil ions is particularly significant for enzyme inactivation and intracellular damage. As shown in Table – 3, the energy deposited by these particles per unit length is sufficient to break various critical biological bonds, including the relatively strong covalent bonds like S-S (Disulfide) and P-O (in ATP/ADP), as well as weaker non-covalent bonds such as Van der Waals and Hydrogen bonds that are crucial for maintaining the three-dimensional structure and function of enzymes and DNA. The recoil of Al-28 and Si-28 ions, although occurring over a very short range (0.022 – 1.5 Å), can directly disrupt the active sites of enzymes due to the momentum transfer and local structural distortion. These signs together with the difference in charge and ionic radius from Mg^2+^ to Al^3+^ to Si^4+^ will certainly inactivate Mg-dependent enzymes once Mg-28 replaces stable Mg in these enzymes. Specifically, the inactivation of DNA/RNA polymerases, helicase, topoisomerase, and DNA repair enzymes disrupts or arrests the S phase of the cell cycle, leading to apoptosis. Targeting telomerase prevents cancer cells from achieving immortality, thus limiting tumor growth and reducing the potential for invasion. Furthermore, the inactivation of matrix metalloproteinases (MMPs) impairs the ability of cancer cells to invade surrounding tissues and metastasize. By disrupting key metabolic enzymes such as hexokinase, kinases, and ATPases, Mg-28 deprives cancer cells of essential energy, leading to metabolic collapse and necrosis.

Beta-minus particles, despite their lower LET, have a longer range that allows them to traverse larger cellular distances, generating free radicals along their path. These free radicals can indirectly damage DNA, proteins, and other cellular components, contributing to the overall cytotoxic effect of Mg-28. The combined action of direct bond breakage by high-LET particles and indirect damage by free radicals ensures a multifaceted attack on essential cellular machinery.

Targeted irradiation within the nucleus and mitochondria is achieved through the selective uptake of Mg-28, where crucial Mg-dependent enzymes like DNA/RNA polymerases, hexokinase, and telomerase reside. The subsequent decay of Mg-28 directly induces localized irradiation via beta particles, Auger electrons, and recoil ions, resulting in a concentrated release of damaging particles close to their molecular targets, thus maximizing enzyme inactivation and cellular destruction.

### IV.4. Precision Targeting and Safety Profile

The distinct ranges of the emitted particles from Mg-28 decay contribute to its precision targeting and favorable safety profile. The very short range of recoil ions (0.022 – 1.5 Å) ensures that their destructive energy is deposited within nanometer distances, primarily affecting the enzyme molecules in their immediate vicinity. Auger electrons, with a slightly longer range (88 – 224 nm), also deposit their energy within the subcellular compartments, causing localized damage to organelles like the nucleus and mitochondria.

In contrast, the longer range of beta-minus particles (up to 6.11 mm) might suggest potential for off-target effects. However, the preferential uptake of Mg-28 by cancer cells, as indicated by the high Mg-uptake coefficient, concentrates the source of these beta particles within the tumor tissue. Furthermore, the energy deposition per unit length (LET) of beta-minus particles is lower compared to Auger electrons and recoil ions, meaning that while they can travel further, the density of ionization events along their path is less intense. This localized delivery of radiation, particularly the high-LET emissions within the tumor cells, minimizes the exposure of surrounding healthy tissues to significant radiation doses.

The absence of a need for bulky biochemical carriers or nanoparticles further enhances the precision and safety of Mg-28 therapy. As a naturally recognized ion, Mg-28 is efficiently transported into cancer cells, ensuring direct intracellular irradiation without the complexities and potential drawbacks of exogenous delivery systems. This intrinsic targeting mechanism contributes to the high local dose within the tumor while minimizing systemic toxicity. These are illustrated on Figure-1.

**Figure 1.**
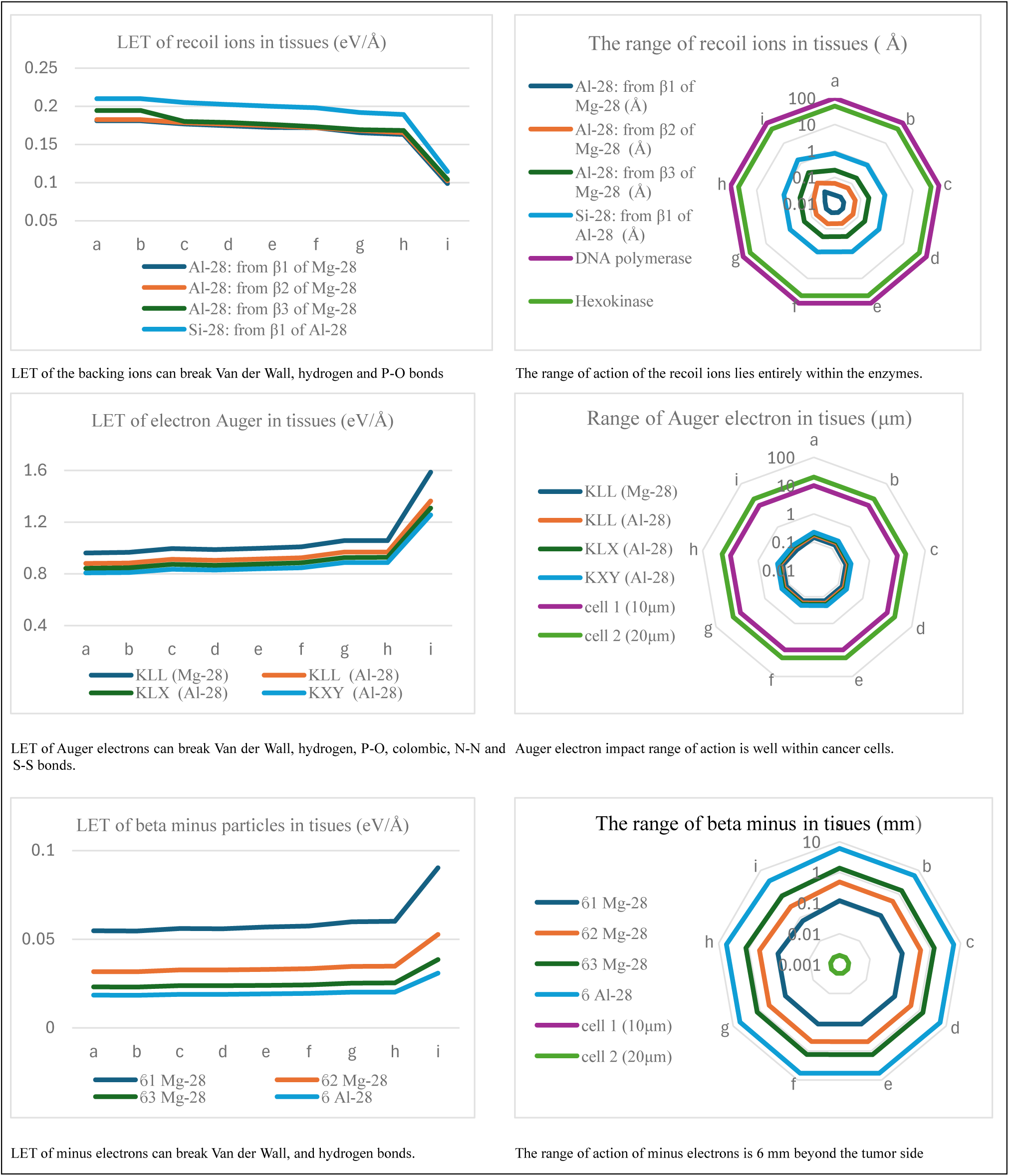
LET and Range of particles in tisues (data derived from. **Table 2 and Table 3)**. Note: a) water; b) Tissue, Soft (ICRP); c) Brain; d) Musco skeletal (ICRP); e) Lung; f) Blood; g) Skin (ICRP); h) M-E Liquid, Sucrose; i) Bone Cortical (ICRP).

### IV.5. High Local Dose with Minimal Systemic Toxicity

Simulation results by MIRD code [28], presented in Table – 4, demonstrate the potential of Mg-28 to deliver a high absorbed dose to tumors while maintaining minimal systemic toxicity. The calculations, performed over a 21-hour period (approximately one half-life of Mg-28), considered different treatment regimens based on the number of Mg-28 ions internalized per cancer cell.

Assuming a regimen of 400 Mg-28 ions per cell (case f), a tumor absorbed dose of 415 Gy can be achieved across all tumor stages (T_0_-T_5_). This dose is significantly above the generally accepted therapeutic threshold of around 50 Gy required for effective tumor control. Notably, the total amount of Mg-28 needed to deliver this high local dose is remarkably small, with a maximum of only 6.2 ng required for even the largest T_5_ tumors. This high efficacy at the nanogram scale underscores the potency and economic viability of the Mg-28 approach.

Conversely, the corresponding whole-body absorbed dose for the same regimen (case f) remains consistently low, with a maximum of 2.33 Gy observed for the smallest T_0_ tumors and decreasing to the mGy range for larger tumors (e.g., 59.2 mGy for T_2_ and 5.92 mGy for T_4_). This significant disparity between the high local tumor dose and the low systemic exposure highlights the inherent safety advantage of Mg-28 therapy, minimizing potential damage to healthy organs and tissues.

The enhanced Mg uptake by cancer cells, driven by their elevated metabolic and replicative demands, further contributes to this favorable dose distribution. This biological preference ensures that Mg-28 is selectively concentrated within the tumor microenvironment, maximizing the therapeutic effect while sparing normal cells.

The data in Table 4 also illustrates the dose-response relationship with varying numbers of Mg-28 ions per cell (cases d, e, and f). Even with lower intracellular concentrations of Mg-28 (62 and 300 ions/cell), therapeutically relevant tumor doses (60.1 Gy and 312 Gy, respectively) can be achieved with correspondingly lower systemic exposure and amounts of Mg-28 required. This flexibility in dosing regimens allows for tailored treatment strategies based on tumor characteristics and patient-specific factors. Figure-2 illustrates these discussions.

**Figure 2.**
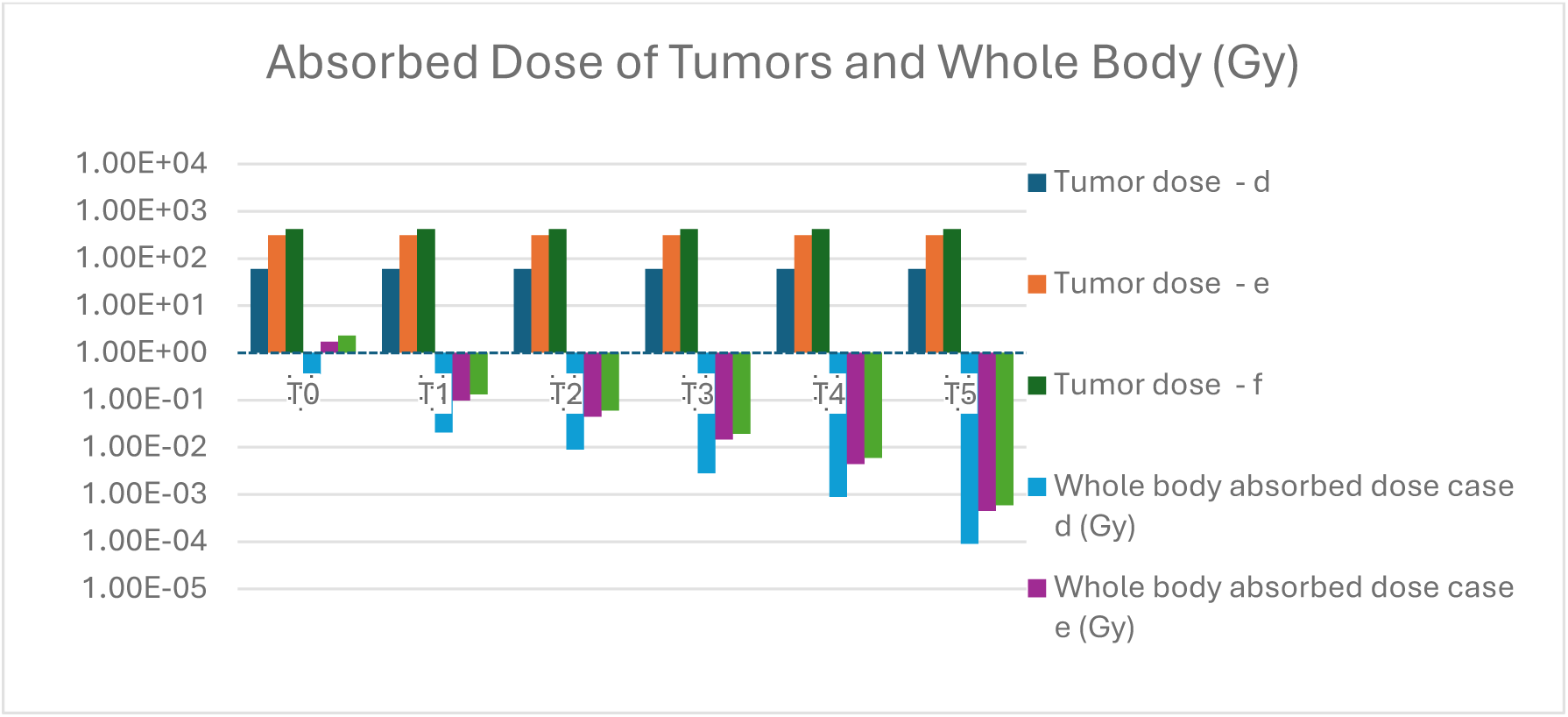
Tumor Absorbed and Whole body Doses in Gy (data derived from. **Table 4)**. *Note: i). In this figure, the tumor absorbed dose and systemic dose are taken from Table 4*. *ii). The absorbed dose to the tumors was calculated based on the assumption that the number of Mg-28 ions taken up by the cancer cells to inactivate Mg-dependent enzymes was: 62, 300 and 400 ions, respectively*. *iii). The systemic absorbed dose was also calculated based on this assumption*. *iv). The absorbed dose at the tumor is in the range of 60-415 Gy, while the whole body dose is in the range of 1.0E-4 Gy to 2.4 Gy. Absolutely safe for the patient but kills most of the cancer cells*.

### IV.6. Summary of Key Advantages

The Mg-28 therapy presents a transformative and biologically intelligent approach to cancer treatment, offering several key advantages that distinguish it from conventional modalities:

a. ***Dual Mechanism of Action for Enhanced Efficacy*:** By simultaneously targeting multiple critical Mg-dependent enzymes essential for cancer cell survival and proliferation (DNA/RNA polymerases, hexokinase, telomerase, MMPs) and delivering highly localized intracellular irradiation through its decay products (beta particles, Auger electrons, and recoil ions), Mg-28 offers a synergistic therapeutic effect that can overcome drug resistance and achieve comprehensive tumor control, a key advantage over single-target traditional inhibitors.
b. ***Intrinsic and Highly Selective Tumor Targeting*:** The therapy leverages the inherent metabolic vulnerability of cancer cells, characterized by their rapid proliferation and consequently elevated demand for magnesium. The quantified Mg-uptake coefficient demonstrates a significant preferential accumulation of Mg-28 within tumor tissues, eliminating the need for complex and potentially toxic biochemical carriers or nanoparticles. This natural targeting mechanism ensures a high concentration of the therapeutic agent directly within the tumor microenvironment.
c. ***Integrated Self-Theranostic Capability*:** Mg-28 uniquely combines diagnostic and therapeutic functionalities within a single radioisotope, embodying a self-theranostic approach. Its gamma emissions allow for early tumor detection using PET or SPECT imaging, even at microscopic stages, while its continuous uptake facilitates real-time monitoring of treatment response, enabling personalized and adaptive therapeutic strategies without the need for additional agents.
d. ***Maximized Local Efficacy with Minimized Systemic Toxicity:*** Simulation data indicate that Mg-28 can deliver cytotoxic radiation doses directly to tumor cells at the nanogram scale, while the absorbed dose to surrounding healthy tissues remains remarkably low. This favorable therapeutic index is attributed to the selective tumor uptake and the short-range, high-LET emissions of its decay products, concentrating the destructive energy within the tumor volume.
e. ***Broad Applicability and Potential for Expanded Therapeutic Horizons:*** The fundamental principle of exploiting the increased metabolic demands of rapidly dividing cells suggests that Mg-28 therapy holds promise across a wide spectrum of cancer types and stages. Furthermore, the concept of targeting host cell enzymes essential for pathogen replication opens avenues for exploring its utility in treating other diseases, such as viral infections, including coronaviruses which rely on host cell RNA polymerase for replication.

## V. Conclusion

In conclusion, the Mg-28 method represents a revolutionary and multifaceted approach to cancer therapy, employing a dual mechanism of action: targeted inactivation of crucial Mg-dependent enzymes and highly localized intracellular irradiation.

The intrinsic selectivity of this method is elegantly driven by the elevated magnesium demand of cancer cells, a phenomenon mathematically quantified by the Mg-uptake coefficient. This biological imperative eliminates the need for complex biochemical carriers or nanoparticles, enhancing both the precision of treatment and its safety profile.

The foundational hypothesis of leveraging Mg-28 in oncology has been supported by preliminary analytical modeling and simulations of LET and absorbed dose, demonstrating the potential for achieving cytotoxic effects at therapeutic thresholds with even nanogram-scale doses. Importantly, the behavior of Mg-28 within the tumor microenvironment embodies a form of natural biological targeting, where cancer cells, fueled by their metabolic and replicative demands, act as selective attractors for Mg²⁺. This pathway allows Mg-28 to efficiently infiltrate intracellular compartments, particularly the nucleus and mitochondria, where it disrupts the enzymatic machinery critical for cancer progression.

Furthermore, the inherent gamma emissions of Mg-28 confer an integrated self-theranostic capability, enabling early tumor detection and real-time monitoring of treatment response, offering unparalleled precision in cancer management. Significantly, this approach holds promise for application across all cancer types and stages, a distinct advantage over many existing therapies that are often limited by tumor type or stage of progression. Beyond oncology, the Mg-28 approach could also be explored for treating viral infections that rely on host RNA polymerase for replication, including coronaviruses, which utilize the host cell’s RNA polymerase for their genome replication. By inactivating this enzyme, Mg-28 may disrupt the viral replication cycle, offering a novel therapeutic strategy. This innovative approach provides a solid foundation for future preclinical (in vitro, in vivo) and clinical investigations to confirm its therapeutic efficacy, pharmacokinetics and biosafety. It deserves further study to develop a new hypothesis in cancer treatment.

## VI. Challenges and Recommendations

**Challenges**:

a. Clinical safety and efficacy require extensive evaluation.
b. Mg-28 production is costly and technically complex.
c. Clinical trials demand significant resources.
d. Necessitates extremely rapid and efficient transportation and utilization procedures of Mg-28.

**Recommendations:**

1. Conduct in vitro and in vivo studies to validate the model.
2. Establish Cancer Hospitals Near Nuclear Facilities.
3. Research and Develop On-Site Mg-28 Production Devices
4. Establish Prioritized Transportation Networks.

## Acknowledgments

Thanks to Dr. Vu Thien Y (Pharmaceutical Faculty, Ton Duc Thang University, Vietnam) for valuable feedback during manuscript preparation.

## Author Contribution

Dr. Tran Van Luyen designed the hypothesis, performed simulations, analyzed data, and wrote the manuscript.

## Ethical Statement

This study adheres to ethical principles in research and data dissemination.

---

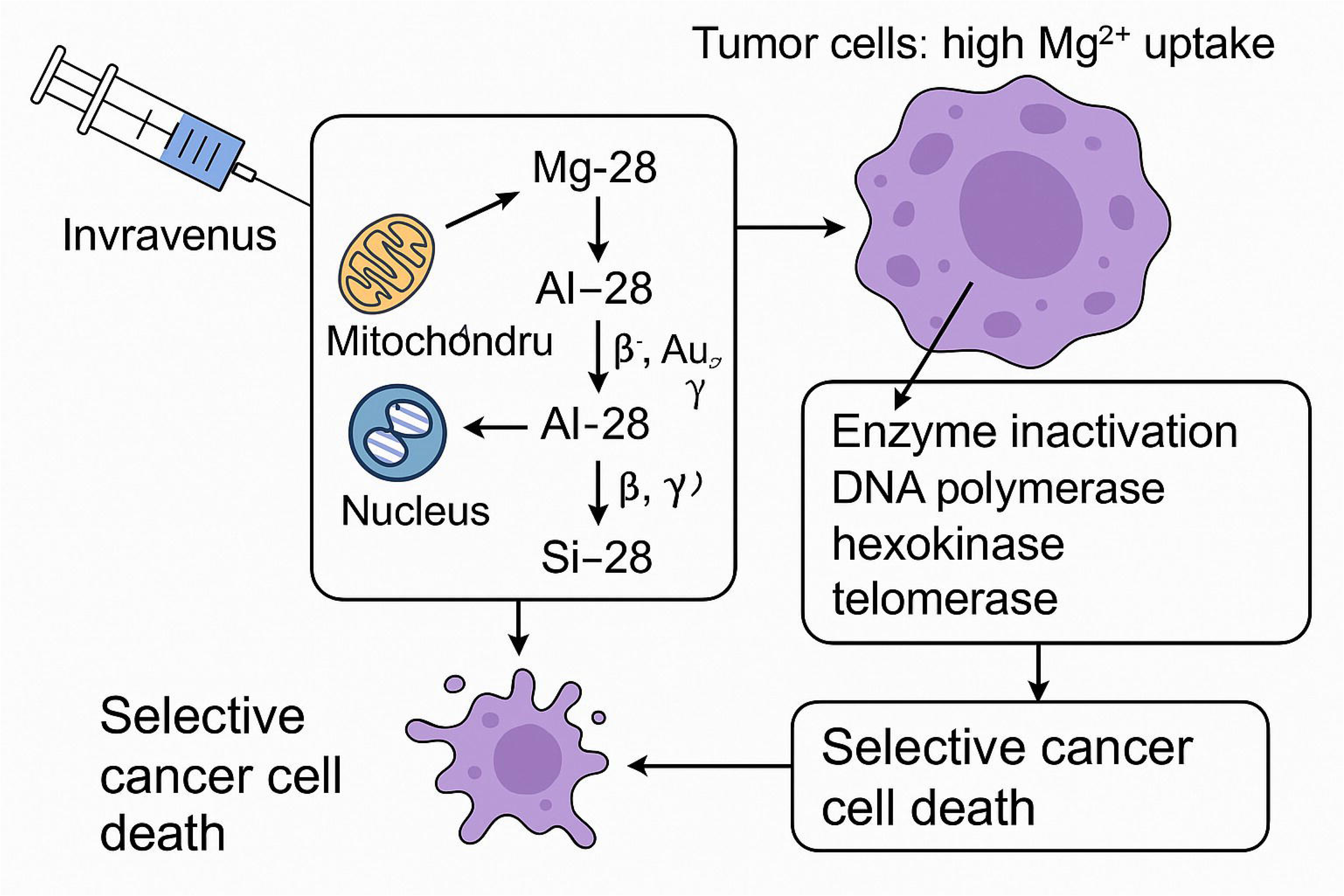

